# Structural basis for CDK7 activation by MAT1 and Cyclin H

**DOI:** 10.1101/2020.08.20.258632

**Authors:** Stefan Peissert, Andreas Schlosser, Rafaela Kendel, Jochen Kuper, Caroline Kisker

**Affiliations:** Rudolf Virchow Center for Integrative and Translational Bioimaging, Institute for Structural Biology, University of Würzburg, 97080 Würzburg, Germany; Comprehensive Cancer Center Mainfranken, Würzburg, Germany

**Author notes:** Corresponding authors: C Kisker and J Kuper. **Author contributions**, The author(s) have made the following declarations about their contributions: Conceived and designed the experiments: SP, JK and CK. Performed the fluorescent based kinase assay: RK. Performed all other experiments except for mass spectrometry and analyzed the data: SP. Performed the mass spectrometry experiments and analyzed the data: AS. Refined the structures: SP and JK. Contributed to the writing of the manuscript: SP, JK and CK.

**Keywords:** CDK-activating kinase, transcription, cell cycle control, kinase

## Abstract

CDK7, Cyclin H, and MAT1 form the heterotrimeric CDK-activating kinase (CAK) complex. CAK is a vital factor for the two essential processes of transcription and cell cycle control. When associated with the general transcription factor II H (TFIIH) it activates RNA polymerase II by hyperphosphorylation of its C-terminal domain (CTD). In the absence of TFIIH it phosphorylates the T-loop of CDKs that control cell cycle progression. CAK holds a special position among the CDK branch due to this dual activity and the dependence on the MAT1 protein for activation in addition to Cyclin H. We solved the structure of the CAK complex from the model organism *C. thermophilum* at 2.6 Å resolution. Our structure reveals an intricate network of interactions between MAT1 and its two binding partners CDK7 and Cyclin H providing a structural basis for the mechanism of CDK7 activation and CAK activity regulation. *In vitro* activity measurements combined with functional mutagenesis show that CDK7 activation can occur independently of T-loop phosphorylation and is thus exclusively MAT1 dependent by positioning the CDK7 T-loop in its active conformation. Finally, our structure of the active CAK with a peptide model provides a molecular rationale for heptad repeat phosphorylation.

**Significance Statement:** The fundamental processes of cell cycle regulation and transcription are linked by the heterotrimeric CDK-activating kinase (CAK) complex. We have solved the crystal structure of the active CAK complex and provide a molecular rationale for CAK activation, regulation, and substrate recognition thereby highly advancing our understanding of this essential factor.

## Introduction

Transcription and cell cycle control are two essential processes which are tightly regulated by cyclin-dependent kinases (CDKs). The activity of CDKs is tightly controlled by two events: (1) Phosphorylation of the T- or activation-loop and (2) through the interaction with their respective cyclins. Structural studies have shed light on the activation and regulation of heterodimeric CDK/Cyclin complexes [reviewed in (Wood and Endicott 2018)]. Interestingly, the cyclins share little sequence homology amongst each other but all contain either one or two versions of the cyclin box (Gibson, Thompson et al. 1994, Malumbres 2014). All CDKs exhibit a central kinase domain and variable N- and C-terminal domains. The vital activation loop is located in between the DFG and an APE motifs within the kinase domain (Wood and Endicott 2018). As exemplified for CDK2, activation usually takes place sequentially through phosphorylation of Thr160 in the activation loop by a CDK-activating kinase (CAK) and subsequent binding to Cyclin A (Merrick, Larochelle et al. 2008).

Amongst the human CDK family CDK7 assumes a unique and highly important position. CDK7 is not only at the direct interface between the important processes of cell cycle regulation and transcription but also, in contrast to other CDKs, requires two proteins for its activation, i.e. Cyclin H and the RING finger protein MAT1. The heterotrimeric complex formed by these proteins activates the cell cycle CDKs 1, 2, 4 and 6 via T-loop phosphorylation (Lolli and Johnson 2005) and in humans is thus referred to as the CAK complex. Compared to the other CDKs, however, formation of the trimeric CAK complex does not necessitate phosphorylation of the threonine in the activation loop to activate the kinase (Devault, Martinez et al. 1995, Fisher, Jin et al. 1995, Tassan, Jaquenoud et al. 1995).

During transcription, CDK7 phosphorylates the C-terminal domain (CTD) of the RNA polymerase II subunit Rpb1 (Feaver, Svejstrup et al. 1994, Roy, Adamczewski et al. 1994, Akoulitchev, Makela et al. 1995, Shiekhattar, Mermelstein et al. 1995) at Ser5 and Ser7 of the CTD heptad repeat YSPTSPS (Komarnitsky, Cho et al. 2000, Akhtar, Heidemann et al. 2009, Glover-Cutter, Larochelle et al. 2009, Kim, Suh et al. 2009) thereby disrupting the interaction between the mediator and the RNA polymerase (Myers, Gustafsson et al. 1998, Sogaard and Svejstrup 2007) and thus triggering the release of RNA polymerase II and enabling promoter clearance. CTD phosphorylation was found to be enhanced when the activation loop threonine Thr170 of CDK7 was phosphorylated *in vitro* by the CDK2/Cyclin A heterodimer (Larochelle, Chen et al. 2001) suggesting a regulatory role towards substrate specificity through this phosphorylation event. Furthermore, CDK7 is involved in promoter proximal pausing, co-transcriptional chromatin modification and termination (Glover-Cutter, Larochelle et al. 2009, Ebmeier, Erickson et al. 2017) emphasizing the essential role during the entire process of RNA Pol II based transcription. Recent studies have also implicated the CAK in the resolution of phase separated RNA Pol II droplets to enable transcription initiation (Boehning, Dugast-Darzacq et al. 2018). During transcription the MAT1 subunit not only activates the CAK but also tethers it to the core of the general transcription factor II H (TFIIH) ensuring the correct positioning of the active complex directly at the target site (Busso, Keriel et al. 2000, Abdulrahman, Iltis et al. 2013). At the same time MAT1 also controls core TFIIH function (Sandrock and Egly 2001, Kokic, Chernev et al. 2019, Peissert, Sauer et al. 2020) further highlighting its essential role for the CAK complex and transcription initiation.

Since a sustained proliferative signaling and the evasion of growth suppressors are both hallmarks of cancer (Hanahan and Weinberg 2011), it is not surprising that CDKs and cyclins are often overexpressed and CDK inhibitory proteins are dysregulated in cancer cells (Bai, Li et al. 2017). The potential of CDKs for the therapeutic intervention in cancer treatment (Sanchez-Martinez, Lallena et al. 2019) has been exploited for CDK7 resulting in the selective CDK7 inhibitor THZ1 (Kwiatkowski, Zhang et al. 2014) and its derivate SY-1365 (Hu, Marineau et al. 2019) that have been proven to specifically eliminate cancer cells that heavily rely on high levels of transcription such as triple negative breast cancer (Wang, Zhang et al. 2015, Li, Ni Chonghaile et al. 2017).

Despite its vital function in cellular processes and implications for cancer, the structural information on the CAK complex was limited to the single subunits Cyclin H and inactive CDK7 (Kim, Chamberlin et al. 1996, Andersen, Busso et al. 1997, Lolli, Lowe et al. 2004) impeding a detailed understanding of CDK7 activation and regulation by its binding partners. Here, we report the crystal structures of the apo and ATPγS bound CAK complex from *C. thermophilum* which disclose the extensive interaction network within the heterotrimeric complex leading to the active form of CDK7. While CDK7 and Cyclin H are engaged in conserved interactions between the N-terminal lobes, MAT1 is embedded in a gap between the C-terminal lobes of CDK7/Cyclin H extensively interacting with both proteins and thereby significantly stabilizing the heterotrimeric complex. The activation loop of CDK7 is tethered in a catalysis-competent conformation by Cyclin H and especially MAT1. Structure based mutagenesis and subsequent functional analysis confirm the unique mechanism by which MAT1 activates CDK7.

## Results

### Biochemical characterization of the CAK complex

Towards the functional and structural characterization of the CAK complex, we recombinantly expressed and purified the homologues of CDK7 (ctCDK7), Cyclin H (ctCyclin H), and MAT1 (ctMAT1) from the model organism *Chaetomium thermophilum*. All proteins share a high degree of sequence conservation with their human and yeast homologues (Supplementary Figure 1) indicating high functional conservation. Full complex formation of the heterotrimeric complex was achieved using a heterologous approach. ctCyclin H and ctMAT1 where co-expressed in *E. coli*, whereas ctCDK7 was expressed in insect cells and purified separately (see Methods). ctMAT1 was expressed either as full-length protein or as the CAK interaction module (Busso, Keriel et al. 2000) containing residues 250-338 (ctMAT1short). Heterotrimeric complex formation was achieved by mixing the purified proteins and subsequent size-exclusion chromatography resulting in the complexes ctCAK or ctCAKshort depending on which MAT1 was used (Fig. 1A, B). Both ctCAK complexes were stable during size exclusion chromatography indicating tight complex formation. SDS PAGE analysis of the purified complexes verified equimolar and homogenous complex formation. To assess the kinase activity properties of the ctCAK complexes we analyzed the phosphorylation activity of ctCAK against a single CTD heptad repeat of the RNA Pol II subunit Rpb1 coupled to GST (GST-YSPTSPS) using an in-gel phospho-staining assay after SDS PAGE (Fig. 1C). CtCAK efficiently phosphorylates the GST-YSPTSPS substrate and the specific phosphorylation of serine 5 in the YSPTSPS motif was confirmed by mass spectrometry (Supplementary Fig. 2 A, B). In contrast, this specific activity was drastically decreased to 8% and 21% for ctCDK7 alone or in the ctCDK7/ctCyclin H heterodimer, respectively (Fig. 1C, bar graph) indicating that both ctCyclin H and ctMAT1 are required for full activation of ctCDK7. Notably this activity was obtained without the requirement of T253 phosphorylation in the activation loop which was obtained almost exclusively unphosphorylated after purification and was not autophosphorylated at T253 during the experiments as confirmed by mass spectrometry (Supplementary Fig. 2 C). Subsequently, we extended our analysis to a more natural substrate and analyzed the kinase activity with the full CTD of the RNA Pol II subunit Rpb1 from *S. cerevisiae* coupled to MBP (scCTD) using a coupled ATPase assay (Fig. 1D). Not surprisingly, this substrate resembling the more natural CTD led to significantly higher activity, however, the relative activation pattern did not change. CtCDK7 by itself is barely active with 5% of ctCAK activity and ctCDK7/ctCyclin H retains 29% of the ctCAK activity showing that the activation requirements do not change between the single heptad peptide and a complete heptad repeat domain. In both experimental setups full activity can only be reached if all three CAK components are present as previously reported (Devault, Martinez et al. 1995, Fisher, Jin et al. 1995, Tassan, Jaquenoud et al. 1995). Importantly, the activity of ctCAKshort is highly comparable to full-length ctCAK (Fig. 1D) confirming that ctMAT1 short is sufficient to activate CDK7 which is well in line with previous data from the Egly group (Busso, Keriel et al. 2000).

**Figure 1:**
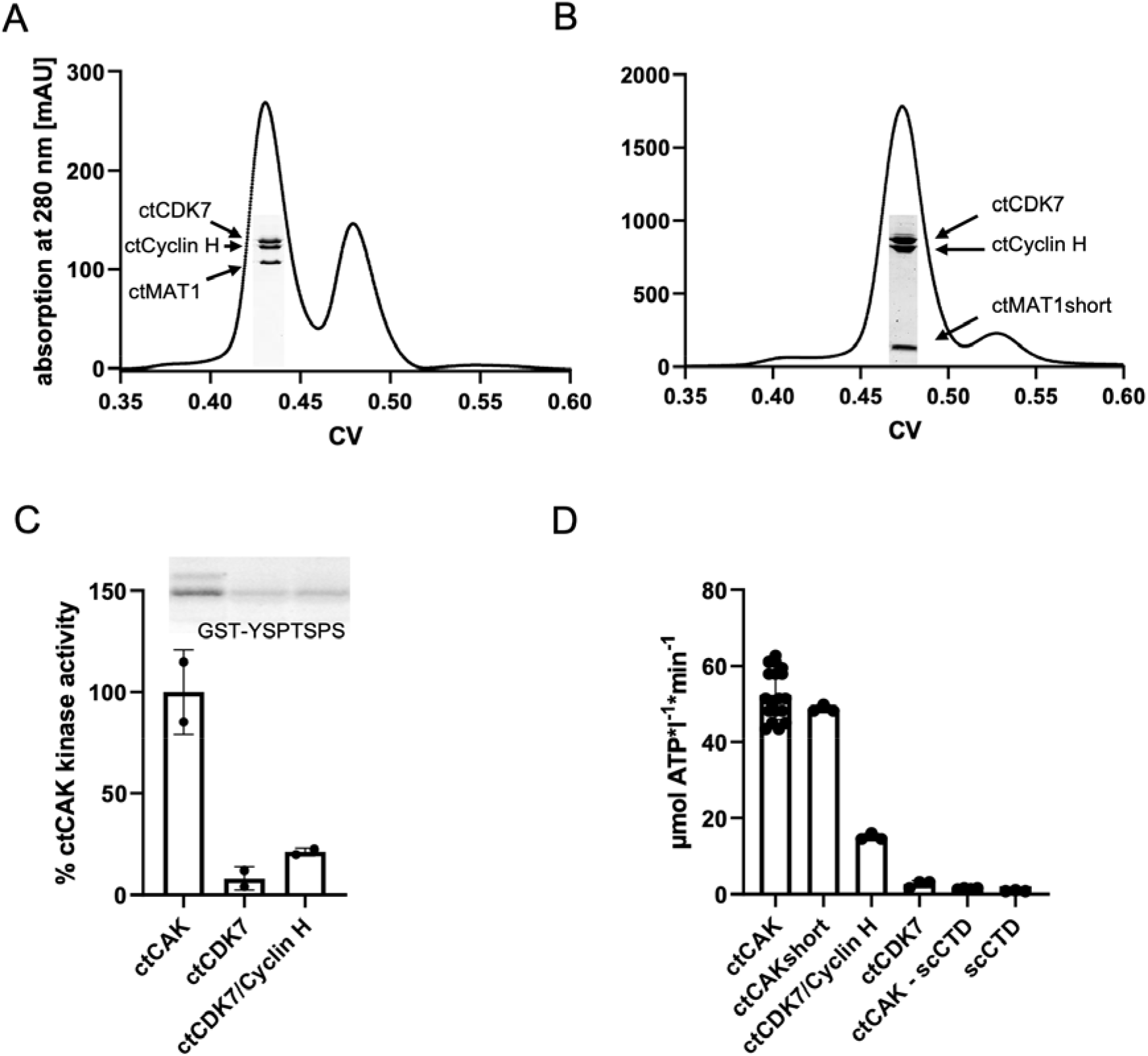
Biochemical characterization of ctCAK complexes. **(A)** Size exclusion chromatography profile of the ctCAK complex. The inset shows the main peak fraction of an SDS PAGE experiment. Proteins are indicated at their respective size. **(B)** As (A) but in this case the formation of ctCAKshort was analyzed. **(C)** Quantification of the phosphorylation activity in the presence of ctCAK, ctCDK7 or ctCDK7/ctCyclin H. Phosphorylation activity was performed with a single heptad peptide coupled to GST and quantified (see methods for details). **(D)** Kinase activity measurements monitoring ATPase function. The indicated proteins were incubated with saturating amounts of ATP and MPB-scCTD substrates in a coupled assay. The highest velocity towards the end of the reaction was fitted and used to determine the ATP consumption over time.

### Overall Structure of the CAK Complex

The ctCAKshort complex was amenable to crystallization and led to the apo structure of the heterotrimeric ctCAKshort complex (Fig. 2A). The structure was solved by molecular replacement using the structures of human Cyclin H and CDK7 as search models (Kim, Chamberlin et al. 1996, Andersen, Busso et al. 1997, Lolli, Lowe et al. 2004). After initial refinement, clear additional difference density permitted *de novo* modeling of ctMAT1. The crystal form belongs to space group P2_1_ and contains two almost identical complexes (rmsd of 0.3 Å) in the asymmetric unit (Supplementary Figure 3 A). In the two heterotrimeric complexes residues 273-338 or 272-338 of ctMAT1 and residues 78-399 or 77-399 of ctCDK7 are resolved in complex A and B, respectively. For ctCyclin H residues 1-4, 50-74, 263-311 and 391-425 or 1-3, 50-74 and 391-425 are disordered in complex A and B, respectively. The final ctCAKshort model was refined to a maximum resolution of 2.6 Å and R-factors of 20.3%/24.9% (R and R_free_, respectively, Table 1).

**Figure 2:**
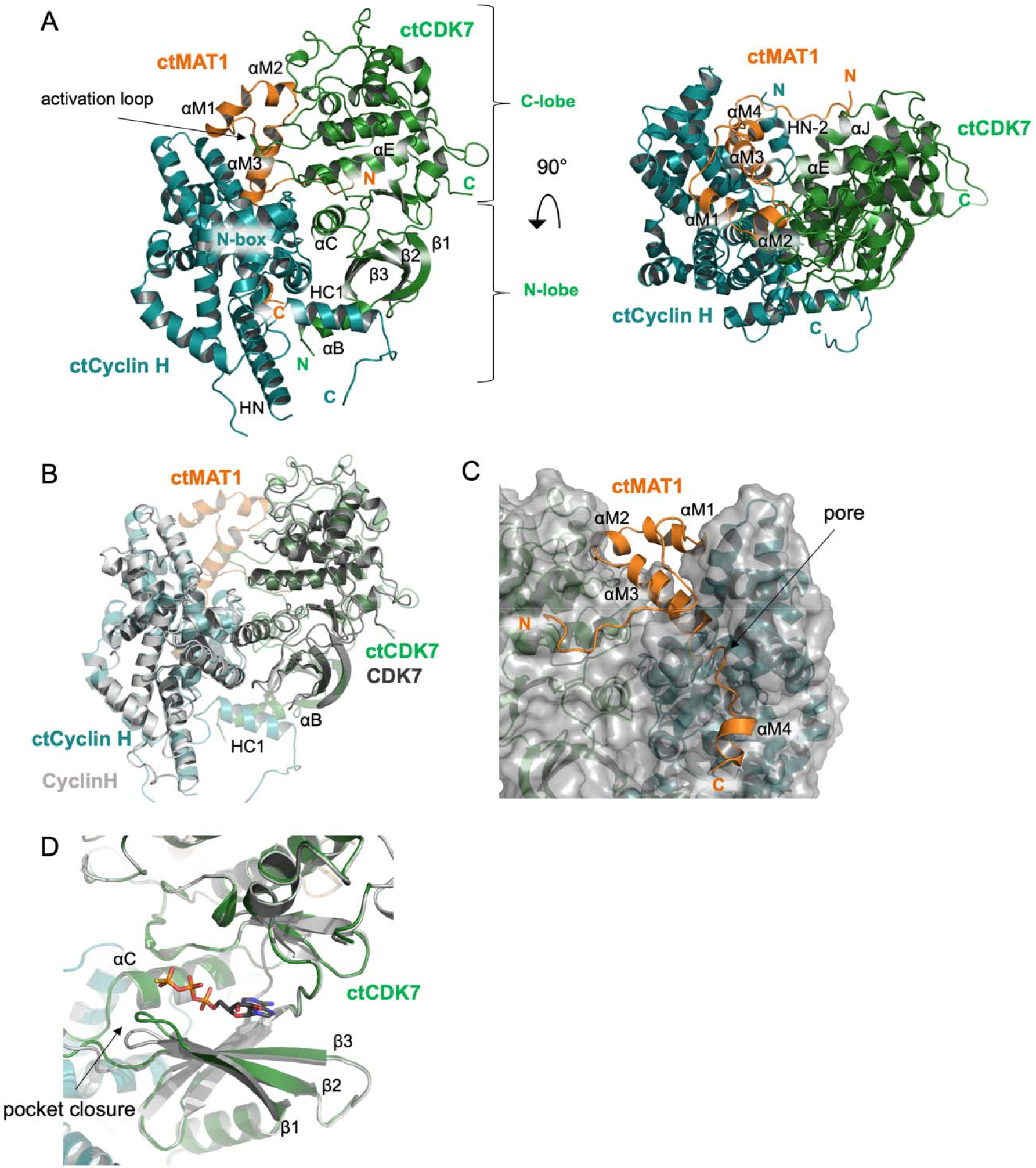
Structure of the ctCAKshort complex from *C. thermophilum*. **(A)** Cartoon representation with two views rotated by 90° of the heterotrimeric ctCAKshort complex. Important secondary structure elements are indicated. CtCDK7 is depicted in green, ctCyclin H in deepteal, and MAT1 in orange. **(B)** Superposition of the human CDK7 (1UA2, dark grey) and Cyclin H (1KXU, light grey) structures onto the ctCAKshort structure. The color coding for ctCAKshort was chosen as in (A). **(C)** The binding interface of ctMAT1 with ctCDK7 and ctCyclin H within ctCAKshort. Color coding as in (A). ctMAT1 is shown in cartoon representation whereas ctCDK7 and ctCyclin H are shown in transparent surface representation with underlying cartoon representation. **(D)** Superposition of the ctCAKapo (grey) and ctCAK ATPγS (green) structures; ATPγS is shown in stick mode. The arrow depicts the pocket closure.

**Table 1.**
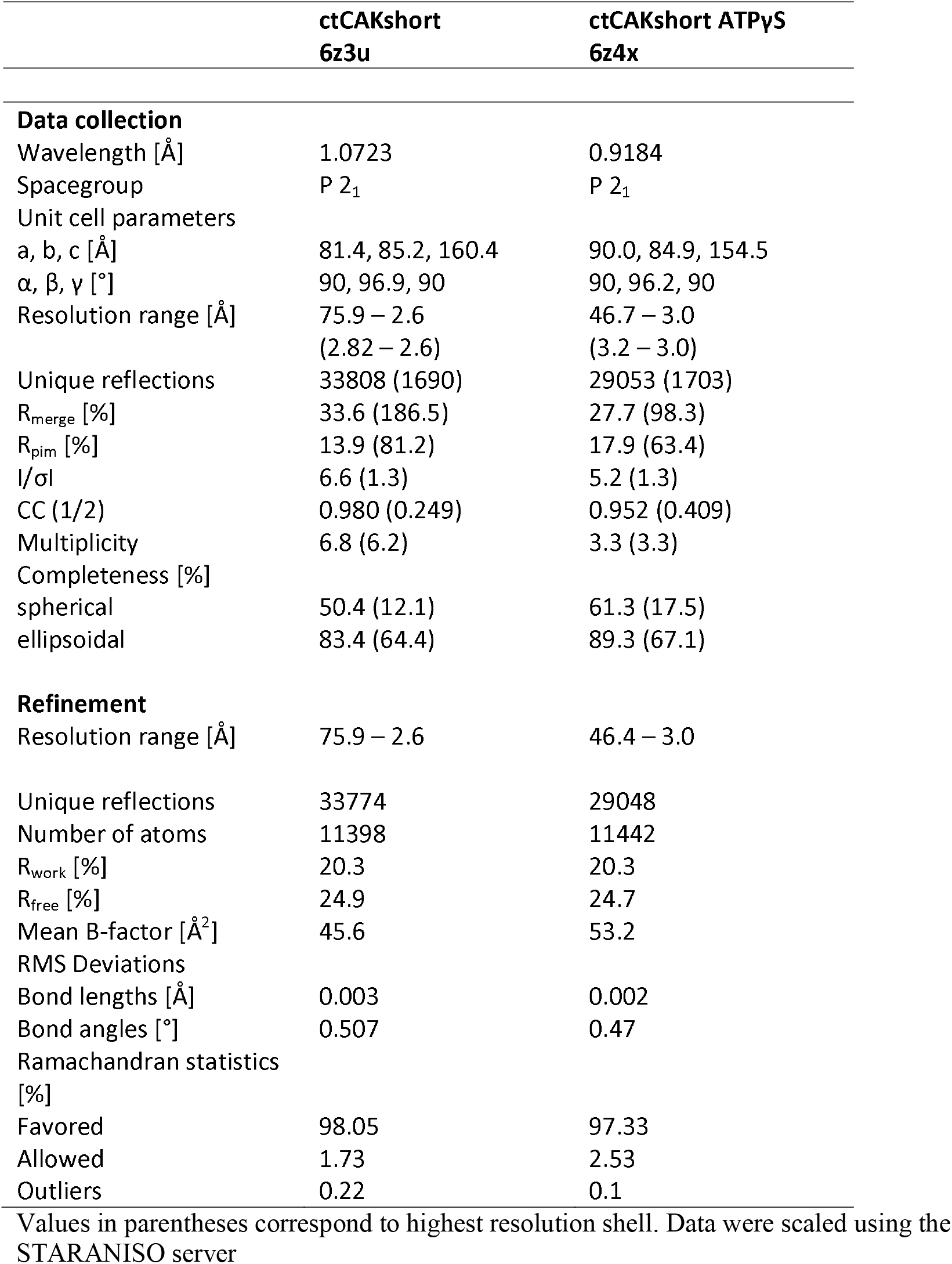
Data collection and refinement statistics

The structure of the heterotrimeric complex reveals a tight interaction of all three subunits with each other. Most contacts between ctCDK7 and ctCyclin H are formed via the N-terminal lobe of ctCDK7 and the N-terminal cyclin box fold of ctCyclin H (Fig. 2A) with a buried surface area of approximately 2950 Å^2^. ctMAT1short adopts a loop rich, extended fold featuring four α-helices in total (Fig. 2A, Supplementary Fig. 1A). All resolved parts of ctMAT1short are essentially involved in protein-protein interactions. Intriguingly, ctMAT1short predominantly seals a large cleft between the C-lobes of ctCyclin H and ctCDK7 thereby contributing a total buried surface area of 5960 Å^2^ thus forming a highly stable complex with extensive interactions. The buried surface area between ctCyclin H and ctMAT1 (3460 Å^2^) exceeds that buried by ctCDK7 and ctMAT1 (2300 Å^2^). However, both ctMAT1 interfaces bury more surface area than through the interaction between ctCDK7 and ctCyclin H indicating the importance of MAT1 for the stability of the CAK complex as previously described (Devault, Martinez et al. 1995, Fisher, Jin et al. 1995, Tassan, Jaquenoud et al. 1995).

The superposition of the individual subunits of the ctCAK complex with human Cyclin H and CDK7 reveals that not only their sequence but also their folds are highly comparable yielding rmsd values of 1.7 and 1.6 Å, respectively, suggesting a high functional conservation (Fig. 2B). A closer look at the CDK7 superposition shows that an additional N-terminal α-helix (αB), is resolved in ctCDK7 which interacts with HN and HC1 of ctCyclin H, of which HC1 was not resolved in the apo structure of human Cyclin H. Another noticeable difference is the loop between β3 and the C-helix of ctCDK7 which is now resolved and could not be observed in the human CDK7 structure. Additionally, an N-terminal extension of ctCyclin H (amino acids 5-20) featuring helices HN-2 and HN-1 can be seen in our ctCAKshort structure which forms extensive contacts with both ctCDK7 and ctMAT1short.

Interestingly, the general architecture of the ctCDK7/ctCyclin H binary complex within the heterotrimeric ctCAKshort structure closely resembles that of other dimeric CDK/Cyclin complexes such as CDK2/Cyclin A, CDK8/Cyclin C and CDK9/Cyclin T (Russo, Jeffrey et al. 1996, Baumli, Lolli et al. 2008, Schneider, Bottcher et al. 2011). One of MAT1’s functions in the trimeric complex is thus to act as a stabilizing factor and tighten the binary complex through its interactions with both CDK7 and Cyclin H.

The N-terminal loop (270-293) of ctMAT1 mostly forms contacts with αE and αJ of ctCDK7 and then traverses the cleft towards ctCyclin H to interact with HN-2. The loop folds back on α-helix αM3 (312-324) of ctMAT1 which is buried in the center of the gap between CDK7 and Cyclin H thereby extensively interacting with both proteins (Fig. 2C). Residues 294-311 of ctMAT1 form αM1 and αM2 that continue back towards ctCDK7 and importantly, form contacts with the activation loop of ctCDK7. αM3 extends towards Cyclin H and the following loop (324-330) closes a pore formed by helices HN-2, HN-1, HN, H2”and H5’ of ctCyclin H as well as ctMAT1 residues 283-286. This pore is formed by the previously unresolved α-helices from Cyclin H that are likely stabilized due to the presence of MAT1 being tightly embedded in the CDK7/Cyclin H interface further emphasizing the special role of MAT1 in the heterotrimeric CAK complex for full complex formation, stabilization and activation. Interestingly the linker in cyclin H between the two helices HN-1 and HN-2 harbors the conserved S15 (S5 in human Cyclin H) which is the primary target of CDK8/Cyclin C mediated inhibition of CAK activity and concomitant blocking of transcription initiation by TFIIH (Akoulitchev, Chuikov et al. 2000).

We also determined the crystal structure of the ctCAKshort complex bound to ATPγS at 3 Å resolution and refined it to R-factors of 20.3%/24.7% (R and Rfree, respectively, Table 1). The conformation of the ATPγS bound ctCAKshort complex is highly comparable to that of the apo structure with an rmsd value of 0.5 Å despite the conformational change of the loop between β-strands β1 and β2 leading to the closure of the ATP binding pocket (Fig. 2D). The nucleotide binding mode of ctCAKshort is similar to that observed for CDK7 and CDK2 (Brown, Noble et al. 1999, Lolli, Lowe et al. 2004).

### Structural basis of CDK7 activation

Our structure provides a first glimpse of CDK7 activation through the formation of the heterotrimeric complex. Two events lead to the active form of the kinase: (1) α-helix H3 of Cyclin H pushes the C-helix of CDK7 towards the active site thus permitting the formation of the salt bridge between the active site lysine K122 (K41 in human CDK7) involved in the phosphotransfer reaction and the conserved glutamate E141 (E62 in human CDK7) from the C-helix. This event also fosters the correct orientation of D238 (D155 in human CDK7) in the DFG motif (‘DFG in’) involved in magnesium binding (Fig. 3A), transforming the kinase to a state capable of ATP hydrolysis. Final activation is achieved by one of the most prominent features revealed in the heterotrimeric ctCAKshort complex: (2) The extensive interaction pattern of the activation loop from ctCDK7 with ctMATshort and partly ctCyclin H leading to a conformation of the activation segment that is highly comparable to the active form of CDK2 (Fig. 3B). The position of the activation loop is stabilized by hydrophobic interactions including the tip of the activation loop around P248 that points into a hydrophobic pocket consisting of residues L296, Y299, Y310, F312 and Y315 from ctMAT1. Two hydrogen bonds between Y310 of ctMAT1 (Y287 in human MAT1) and the carbonyl of A246 of ctCDK7 (G173 in human CDK7) as well as between R190 of ctCyclin H (R165 in human Cyclin H) and D247 of ctCDK7 (S165 in human CDK7) further stabilize the position of the activation loop (Fig. 3C). This intricate network is likely holding the loop in a catalysis-competent position thereby activating ctCDK7 (Fig. 3B, C). Importantly, our structure represents the unphosphorylated state of the activation loop, and thus suggests that this activation mechanism does not require the additional phosphorylation of T253 (T170 in human CDK7) which is well in line with our biochemical data (Fig. 1). However, at the position of T253 we observe that the hydroxy group points towards a positively charged pocket formed by R140 from the re-positioned C-helix, R219 from the HRD motif, and R243 within the activation loop (R61, R163 and K160 in human CDK7, respectively) that seems to be well suited to accommodate the potentially phosphorylated T253 (Fig. 3D). In our structure, this position is occupied by a glutamate from the C-terminus of a symmetry related ctCyclin H subunit (Supplementary Figure 3B) indicating that the complex might be stabilized further once phosphorylation has occurred.

**Figure 3:**
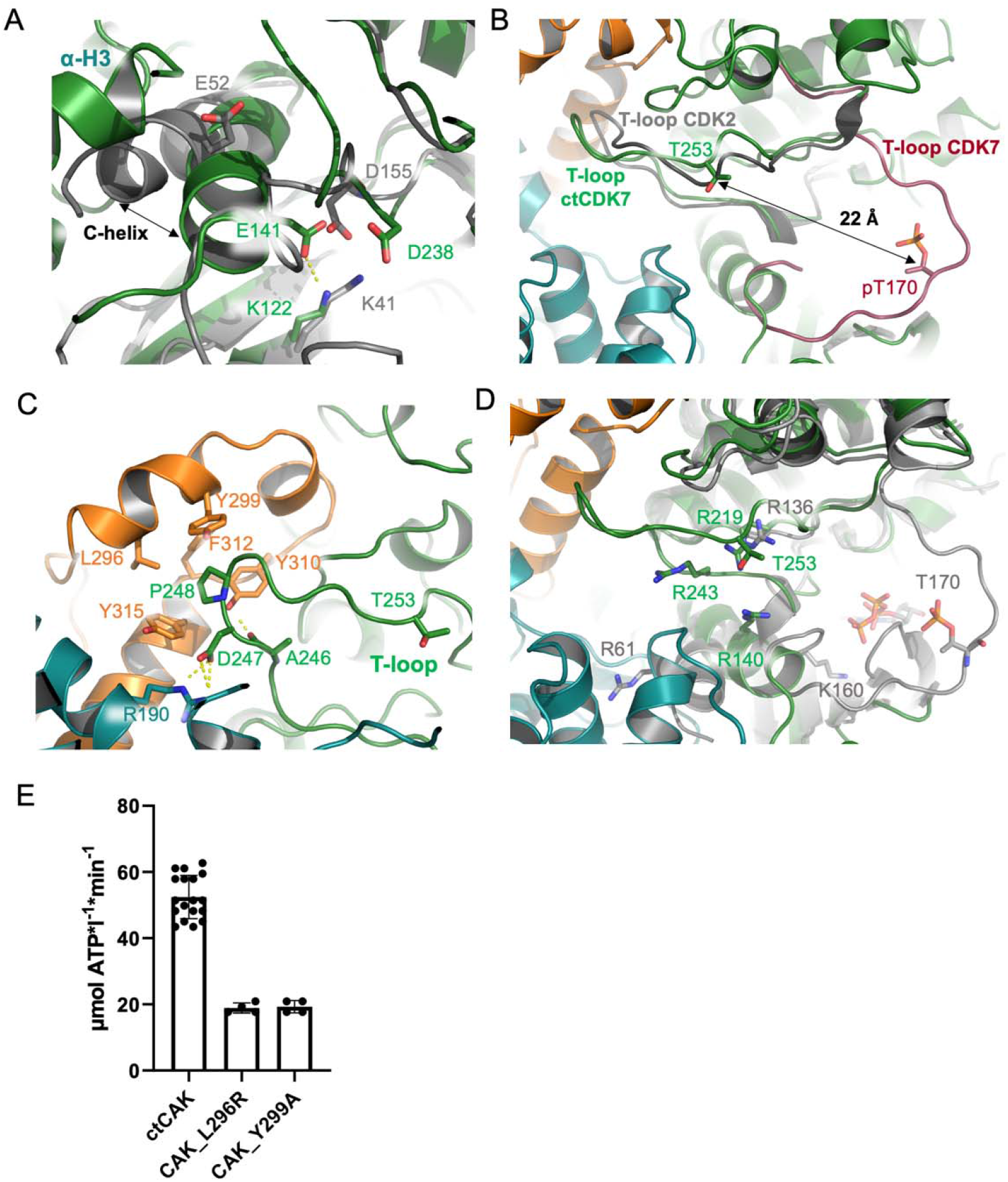
Structural basis of ctCDK7 activation. **(A)** Closeup of a superposition containing human CDK7 (grey, 1UA2) and ctCAKshort (CDK7 in green and Cyclin H in deepteal). The C-helix of CDK7 and relevant active site residues are labeled. **(B)** T-loop movement between ctCAKshort (colored as in (A) and MAT1 in orange), CDK2 (1QMZ, grey) and human CDK7 (1UA2, magenta). The structures were superimposed based on ctCKD7. For clarity only the T-loops of CDK2 and human CDK7 are shown. The T-loop threonine is highlighted. **(C)** ctCDK7 T-loop stabilization via ctMAT1 and ctCyclin H in ctCAK short. Coloring is as in (B). Relevant residues that contribute to the stabilization are shown in ball and stick mode. **(D)** The T-loop phospho-threonine binding site in ctCAK. Closeup of a superposition between ctCAK (based on ctCDK7) and human CDK7 (1UA2, in grey). Coloring is as in (B). Residues are shown in stick mode, labeled and colored according to their respective protein (green CDK7 from the ctCAK complex, grey human CDK7). **(E)** ATPase activity measurements of ctCAK in comparison to ctCAK MAT1 variants (for reference see Figure 1 and the methods section)

To validate our structural observations that the stabilization of the activation loop is MAT1 dependent, we performed structure-based mutagenesis on ctMAT1. The variants were designed to specifically weaken the interaction with the activation loop and not to interfere with the overall interaction pattern. We chose L296 and Y299 (L272 and V275 in human MAT1), with the first residue being strictly conserved and the second type conserved (Supplementary Figure 1) and generated the L296R and Y299A variants. Both variants were used for the assembly of ctCAK complexes as described earlier and subsequently analyzed for their activity by monitoring the phosphorylation of the scCTD substrate. Both, the L296R and the Y299A ctCAK variants showed a significant decrease in kinase activity with 19 μmol ATP l min compared to the wild-type with 52 μmol ATP l^−1^ min^−1^ (Fig. 3E). The reduced activity of the ctCAK variants is similar to that observed for the binary ctCDK7/ctCyclin H complex with 15 μmol ATP l min indicating a complete loss of MAT1 activation (compare Fig. 1D and Fig. 3E). Analysis of the ctCAK complexes containing the MAT1 variants by thermal shift assays showed a decreased melting temperature for both variants suggesting reduced complex stability due to missing contacts in the interface (Supplementary Fig. 4). Importantly, the melting curve is still highly cooperative indicating a single unfolding event of the still intact heterotrimeric complex. Our data therefore suggest that both L296 and Y299 play a significant role towards trimeric complex formation and correct positioning of the CDK7 activation loop.

## Discussion

The human CAK complex assumes a special position among the CDK family due to the dual activity in both transcription and cell cycle progression and it is thus of paramount interest to elucidate the underlying structural activation mechanisms. We solved the crystal structure of the CAK complex from *C. thermophilum* in its active form and shed light on how MAT1 achieves CDK7 activation in cooperation with Cyclin H. The most prominent feature of the ctCAK complex is the intricate interaction pattern of ctMAT1 with its two binding partners resulting in a tightly packed heterotrimeric complex (Fig. 2A) thereby explaining the observed stabilization effect of ctMAT1 on the ctCDK7/Cyclin H heterodimer (Devault, Martinez et al. 1995, Fisher, Jin et al. 1995, Tassan, Jaquenoud et al. 1995). Our structure reveals the two key events which lead to the activation of ctCDK7: (1) α-helix H3 of Cyclin H pushes the C-helix of ctCDK7 towards the active site and (2) the activation loop is stabilized by a hydrophobic pocket formed by residues from ctMAT1 without prior phosphorylation (Fig. 3 A-D). Site directed mutagenesis on this hydrophobic pocket confirmed its importance for CDK7 activation (Fig. 3 E). To our knowledge, the specific ctCDK7 activation mechanism has not been observed for any other CDK/Cyclin pair [reviewed in (Wood and Endicott 2018)] so far. Notably, the other CDK/Cyclin pairs require phosphorylation for activation, whereas ctCAK displays significant activity without this modification. The PHO85/PHO80 complex from *S. cerevisiae* (Huang, Ferrin-O’Connell et al. 2007) and the complex of the Ringo activator Spy1/CDK2 (McGrath, Fifield et al. 2017) can bypass the necessity for phosphorylation like the CAK complex, however, in both cases the binding partner of the kinase mimics the phosphorylation of the activation loop by providing negatively charged residues thus suggesting an entirely different mechanism.

The CDK7/Cyclin H pair within the CAK complex is highly comparable to the overall conformation of other known CDK/Cyclin structures with the conserved contacts between the N-terminal lobes of both proteins (Fig. 4A). Notably, there is some variance in the nature and extent of the interfaces resulting in markedly different rotational angles of the Cyclins with respect to the CDKs, especially when the cell cycle CDKs are compared to the transcriptional CDKs (Fig. 4A). The CDK7/Cyclin H conformation is in between the ‘closed’ form of CDK2/Cyclin A and the ‘open’ form of CDK9/Cyclin T. Another key difference between cell cycle and transcriptional CDKs is the α-helix HN of the Cyclins that forms extensive interactions with cell cycle CDKs but not with transcriptional CDKs. Importantly, α-helix HN in CDK2/Cyclin A and CDK1/Cyclin B (Brown, Korolchuk et al. 2015) is located in a similar position as α-helix M3 of MAT1 suggesting a conserved regulatory mechanism (Fig. 4A). The additionally resolved N-terminal α-helix (αB) of ctCDK7 is comparable to that observed for CDK8 which was described as unique Cyclin C specificity helix (Schneider, Bottcher et al. 2011). Surprisingly, D79 and E82 from the αB-helix of CDK7 form multiple hydrogen bonds with R42, R365 and R375 of ctCyclin H, indicating that despite low sequence conservation among the CDKs and cyclins the function is preserved (Fig. 4A).

**Figure 4:**
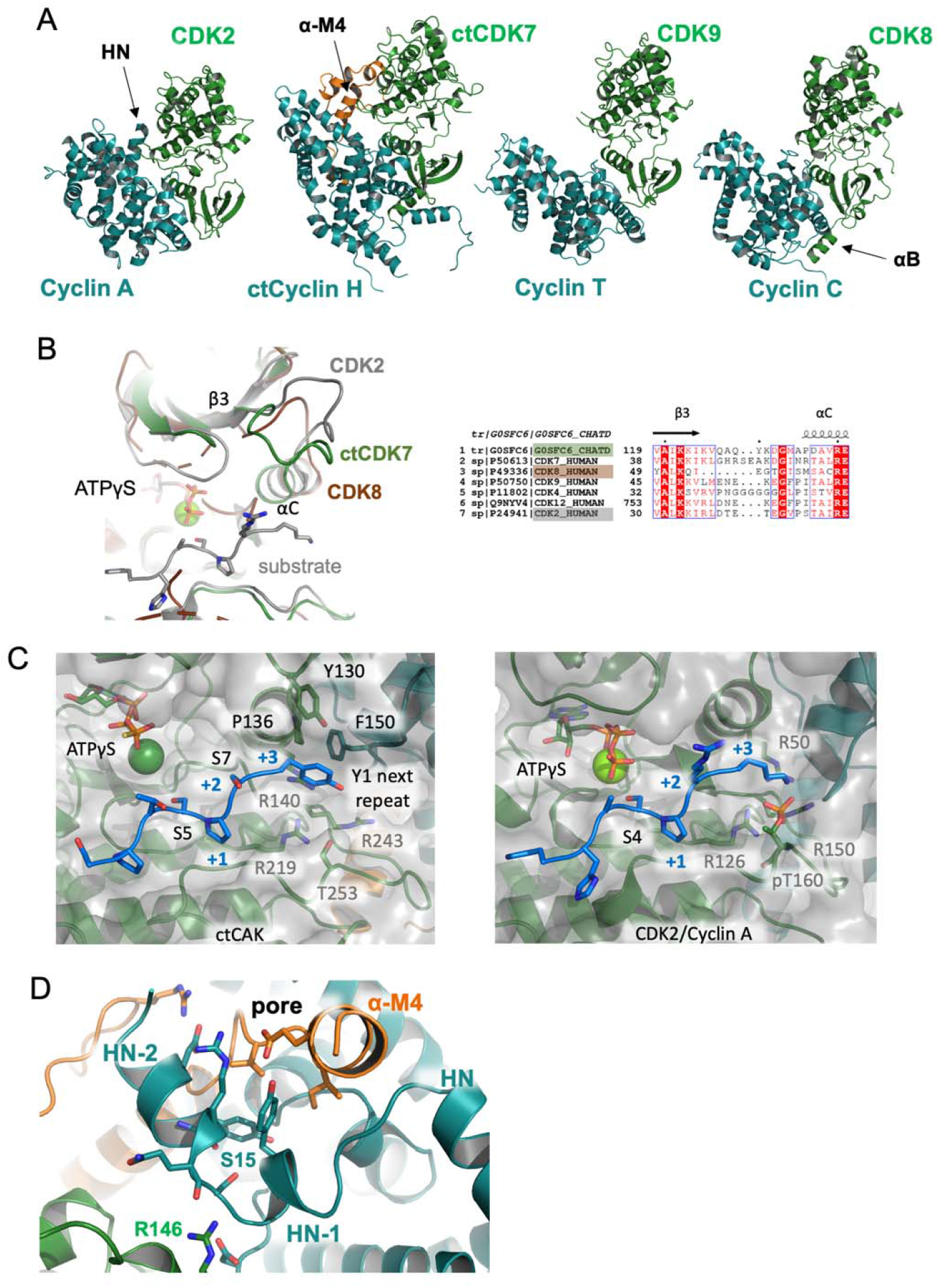
Comparison of ctCAK with other CDK/Cyclin complexes and Substrate Binding. **(A)** Side by side view of ctCAK with structures of CDK2/Cyclin A (1QMZ), CDK9/Cyclin T (3BLH), and CDK8/ Cyclin C (4F7S) based on their superposition. The cyclins are colored in deepteal, the kinases in green. MAT1 is shown in orange. **(B)** Orientation of the loop between γ3 and αC as observed in ctCDK7 (green), CDK2 (1QMZ, grey), and CDK8 (3RGF, brown). The peptide substrate (grey) and the ATPγS are derived from CDK2 and additionally depicted in ball and stick mode. The sequence alignment below the Figure highlights the relevant loop section of ctCDK7 (G0SFC6_CHATD) with other CDKs indicated by their Uniprot entry. The superposed CDKs are highlighted. **(C)** Side by side view of the substrate binding area of ctCAK (left panel) and CDK2/Cyclin A (right panel). CDK is colored green and the Cyclin in deepteal for both panels. Structures are shown in cartoon mode with a transparent surface and specific residues are shown in ball and stick. The peptide model (left panel) and the peptide (right panel) are colored in blue. Relevant positions and residues are indicated. **(D)** Closeup view on a regulatory phosphorylation site in Cyclin H from ctCAK. Color scheme is similar to (A).

Another interesting feature of the CAK complex is the loop between β3 and the C-helix of CDK7 that was not resolved in the inactive human CDK7 structure. Superposition of three exemplary CDK/Cyclin structures indicates that this loop could contribute to the active site cleft formation (Fig. 4 B). The sequence conservation between different CDKs is rather low in this area (Fig. 4B) and it is thus tempting to speculate that the loop between the conserved AXK motif and the C-helix may contribute to substrate binding or specificity.

Our biochemical studies of the ctCAK complex have shown that it specifically phosphorylates a model heptad of the RNA Pol II subunit at position 5 (S5) which is well in line with existing data (Komarnitsky, Cho et al. 2000, Akhtar, Heidemann et al. 2009, Glover-Cutter, Larochelle et al. 2009), and that activation of ctCDK7 requires the presence of ctCyclin H and ctMAT1 (Fig.1 C, D) but importantly does not necessitate T-loop phosphorylation (Supplementary Figure 2). Our data thus implicate that the active site of CDK7, as observed in our structure, resembles an architecture of a catalytically fully competent CDK7. A superposition of our ATPγS bound CAK structure with the active CDK2/Cyclin A structure containing a nucleotide analog and a model substrate peptide shows that the overall architecture of both active sites is well conserved (Fig. 4C, side by side view). This similarity permits a closer look at how CDK7 might recognize the heptad repeat substrate and is able to phosphorylate the S5 position. Based on the CDK2/Cyclin A structure we built a model of the heptad repeat CTD and placed it in our ATPγS CAKshort structure (Fig. 4C, left panel). Intriguingly, if we assume that the S4 of the model substrate assumes the role of S5 in the heptad repeat, this residue would be ideally positioned for the phosphotransfer reaction. The +1 site would be occupied by the vital proline as in the CDK2/Cyclin A peptide preserving the special proline geometry at this position. The next residue in the +2 site would be S7 at the end of the heptad repeat. In the model substrate of the CDK2/Cyclin A structure the following +3 position is a lysine residue that interacts with the phospho-threonine of the T-loop which is additionally stabilized by a positive patch generated by three conserved arginines (Fig. 4C, right panel). Strikingly this position would be occupied by Y1 of the next heptad repeat in our model of the ctCDK7 active site. This next repeat tyrosine would be stabilized by a hydrophobic site created by Y130 and P136 of ctCDK7 and F150 of ctCyclin H. The hydrophobic nature of the site is also conserved in the human and yeast proteins (Supplementary Fig. 1) thus supporting our model. Notably phosphorylation of the T-loop threonine would be well in line with this model since the tyrosine could readily engage in hydrogen bonding to the phospho-threonine as the corresponding lysine does in the CDK2/Cyclin A structure. The surrounding arginine pocket is also well conserved in human CDK7 (Fig. 3D, Supplementary Fig. 1A). Our model thus provides a structural rationale why the functional unit of the heptad substrate is not restricted to one repeat but includes 4 additional residues from the next repeat (Eick and Geyer 2013). The model of substrate processing is also highly compatible with the observation that the CAK could be enhanced/stabilized by T-loop phosphorylation at position 253 (T170 in human CDK7, (Devault, Martinez et al. 1995, Fisher, Jin et al. 1995, Tassan, Jaquenoud et al. 1995, Larochelle, Chen et al. 2001)).

Finally, the CAK complex is not only activated but it can also be inactivated through phosphorylation of S5 and S304 in Cyclin H by the kinase module CDK8/Cyclin C of the mediator complex (Akoulitchev, Chuikov et al. 2000). S5 phosphorylation displayed a more severe phenotype than S304 phosphorylation and was linked to a substantial reduction of CAK activity towards the CTD of RNA Pol II and consequently, deficiencies in transcription initiation. The corresponding residue in ctCyclin H is S15 which is located in a three amino acid long linker between α-helices HN-2 and HN-1 being involved in the described pore (Fig. 4 D). Phosphorylation of this residue requires a change in the local conformation leading to a disruption of the interaction with the C-helix of ctCDK7 and particularly, the C-terminus of ctMAT1short and thus, most likely to complex disruption and thereby to the observed inactivation.

In conclusion, our structure of the fully functional CAK complex provides the long awaited first view on the CDK7 activation mechanism through the tight interaction of CDK7, Cyclin H and the MAT1 revealing the active CAK conformation that is essential for transcription and cell cycle control.

## Materials and Methods

### Molecular biology

For the expression construct of full length ctCyclin H (UniProtKB G0SH78) and ctMAT1 (UniProtKB G0SF48) a modified pET-22 vector (EMBL) was used in which the pelB leader sequence was replaced by the sequence coding for a Twin-Strep-tag (IBA™) and a 3C protease cleavage site (pETtwin, provided by Florian Sauer). The coding sequences of ctCyclin H followed by a second ribosomal binding site and thioredoxinA-6xHis-3C-ctMAT1 were cloned behind the 3C cleavage site of pETtwin by sequence and ligation independent cloning (Li and Elledge 2012). The design of ctMAT1short (250-338) was guided by pairwise protein structure prediction (Yang and Zhang 2015). For the expression constructs of ctCyclin H and ctMAT1short the corresponding coding sequences were inserted behind the 3C cleavage site of pETtwin (ctMAT1short) or pCOLA-SMT3 (ctCyclin H) which is a modified pCOLA vector (provided by Florian Sauer) containing the expression cassette of pETM-22 (EMBL) except that the thioredoxinA-6xHis tag was replaced by a 6xHis-SMT3 tag. The coding sequence corresponding to a single heptad repeat (YSPTSPS) was fused to the C-terminus of GST in a pBADM-30 vector (EMBL). The coding sequence of the RNAP Pol II subunit Rpb1 CTD (UniProtKB P04050, 1534-1733) was amplified from *S. cerevisiae* S288c genomic DNA and cloned into a pETM-41 vector (EMBL) by restriction cloning using NcoI and KpnI resulting in an MBP-scCTD construct.

For the expression in insect cells ctCDK7 was first inserted into the pETtwin vector by SLIC and the resulting Twin-Strep-tag-3C-ctCDK7 construct was then transferred into a pFastBac1 transfer vector (Invitrogen) by restriction cloning using EcoRI and XbaI. The pFastBac1 construct was integrated into the EMBacY baculovirus genome via Tn7 transposition. Mutants were generated according to the Quick-Change site-directed mutagenesis protocol (Stratagene).

### Protein expression and purification

All constructs except for ctCDK7 were expressed in BL21star (DE3) cells (Invitrogen) carrying the pRARE2 plasmid (Novagen) and initially purified using the same protocol. Cells were grown in LB medium supplemented with 100 μg/ml ampicillin (pETtwin) or 50 μg/ml kanamycin (pCOLA-SMT3) and 34 μg/ml chloramphenicol at 37 °C until they reached an OD_600_ of 0.7. The temperature was then reduced to 18 °C and expression was induced with 0.5 mM IPTG overnight. The cells were harvested and resuspended in lysis buffer A (20 mM HEPES pH 7.5, 300 mM NaCl, 25 mM imidazole, 1 mM TCEP) supplemented with 0.5 mM PMSF and DNaseI. After cell disruption the cleared lysate was loaded on a 5 ml HisTrap FF column (GE Healthcare) equilibrated in lysis buffer A and the protein was eluted with a gradient of 25 – 250 mM imidazole. CtCyclin H was incubated with 3C protease overnight at 4 °C to cleave off the Twin-Strep-tag. CtCyclin H, MBP-scCTD or GST-YSPTSPS were concentrated and then loaded on a Superdex 200 16/60 pg column (GE Healthcare) equilibrated in 20 mM HEPES pH 7.5, 250 mM NaCl, 1 mM TCEP. Fractions containing the target protein were concentrated, flash frozen in liquid nitrogen and stored at −80 °C. The co-expressed complex of ctCyclin H and ctMAT1short was loaded on a 5 ml StrepTrap HP column (GE Healthcare) equilibrated in lysis buffer B (20 mM HEPES pH 7.5, 300 mM NaCl, 1 mM TCEP) as a second affinity purification step and eluted with 2.5 mM desthiobiotin. The complex containing fractions were concentrated, incubated with 3C protease and then further processed by size exclusion chromatography as described above. Purification of the full length ctCyclin H/ctMAT1 complex was performed as described for ctCyclin H/ctMAT1short except that buffers additionally contained 10% glycerol.

For the expression of ctCDK7, Sf21 cells (*Spodoptera frugiperda*) were transfected with bacmid DNA using SuperFect (Qiagen) transfection reagent according to the manufacturer’s instructions. The baculovirus was amplified in Sf21 cells before infecting Hi5 cells (*Trichplusia ni*) seeded to 0.5*10^6^ cells/ml with 0.1 V virus titer for an expression for 72 h to express ctCDK7. The cells were harvested, resuspended in lysis buffer B supplemented with 0.5 mM protease inhibitor PMSF and DNaseI. The cleared lysate was further processed by StrepTrap affinity chromatography and size exclusion chromatography as described above.

To assemble the trimeric ctCAK variants, purified proteins were mixed in a stoichiometric ratio and loaded on a SD200 Increase 10/300 GL column (GE Healthcare) equilibrated in 20 mM HEPES pH 7.5, 200 mM NaCl, 1 mM TCEP. Appropriate fractions were concentrated, flash frozen in liquid nitrogen and stored at −80 °C.

### Crystallization, data collection and structure determination

Crystallization trials were set up with the Honeybee 963 robot using the sitting drop vapor-diffusion method. The ctCAKshort complex was crystallized at a protein concentration of 4 mg/ml and with a precipitant containing 100 mM HEPES pH 7.0-8.0, 0.2 M sodium formate, 15-22% PEG 3350. The crystals were flash frozen in the precipitant solution containing in addition 25% ethylene glycol as cryo-protectant.

For the apo structure, data were collected at 100 K at the ESRF beamline ID29. The data were integrated and scaled using XDS (Kabsch 2010) and then merged using the STARANISO server to 2.6 Å (Table 1). The structure was solved by molecular replacement using PHASER (McCoy, Grosse-Kunstleve et al. 2007) and a homology model of ctCyclin H and ctCDK7 obtained from the SWISS-MODEL server (Waterhouse, Bertoni et al. 2018). Two copies of ctCDK7 and ctCyclin H were found in the asymmetric unit. The missing MAT1 was built manually using coot (Emsley, Lohkamp et al. 2010). The structure was improved by alternating cycles of manual model building and automated refinement. The program Buster was used for the initial refinement steps (Bricogne G. and Roversi P 2017) and the final rounds of refinement were performed using the phenix suite (Adams, Afonine et al. 2010). In all refinement steps the anisotropically corrected data were used to the recommended resolution cutoff of 2.6 Å.

For the ATPγS bound structure, crystals were soaked for 2 h in the precipitant solution supplemented with 1 mM ATPγS, 5 mM MgCl_2_ and 1 mM of phosphorylated peptide (YphoSPTSPS) which was not resolved in the structure. Diffraction data of the ATPγS bound crystals were collected at 100 K at the BESSY beamline MX14-1. The data were integrated, scaled and merged to 3 Å as described above. The structure was solved by molecular replacement using the structure of the apo ctCAKshort complex as search model. The structure was refined using Phenix and the anisotropically corrected data to 3 Å. Structural figures were generated using PyMOL.

### *In vitro* gel-based kinase assay

Kinase activity was assessed using a fluorescent in-gel detection of phosphoprotein after SDS-PAGE of the kinase reaction samples. The reactions were carried out in a solution containing 20 mM HEPES pH 7.5, 200 mM NaCl, 1 mM TCEP, 5 mM MgCl_2_, 2 mM ATP and 0.5 μM ctCDK7 or ctCDK7 containing complexes and 2 μM of GST-YSPTSPS substrate. Samples were incubated at 37 °C for 1 h and then, 5 μl of the reaction mix was loaded on a 15% SDS-gel. After gel electrophoresis, the gel was stained for phosphoprotein species using the Pro-Q Phosphoprotein Gel Stain according to the manufacturer’s instructions (ThermoFisher). Signals were detected at an excitation wavelength of 532 nm and an emission wavelength of 605 nm using a PharosFX (BioRad) fluorescent scanner. Signal intensities were quantified using ImageJ (Schneider, Rasband et al. 2012). All measurements were carried out in duplicates and mean values are plotted with their associated SD.

### *In vitro* substrate dependent ATPase assay

Kinase activity was determined using an *in vitro* ATPase assay in which ATP consumption is coupled to the oxidation of NADH via pyruvate kinase and lactate dehydrogenase activities. Kinase activities were assessed at 37 °C, containing 1.5 U pyruvate kinase, 1.9 U lactate dehydrogenase, 2 mM phosphoenolpyruvate, and 0.3 mM NADH, 20 mM HEPES pH 7.5, 200 mM NaCl, 1 mM TCEP and 5 mM MgCl_2_. The reactions were performed with 200 nM of ctCDK7 or ctCDK7 containing complexes and 20 μM of MBP-scCTD as substrate. The mix including all above mentioned components was preincubated at 37 °C until a stable base line was achieved. The reaction was then started by the addition of ATP at a concentration of 1 mM. Activity profiles were recorded at 340 nM using a FluoroStar Optima (BMG Labtech) plate reader until NADH was entirely consumed. The velocity increased during the reaction and the highest velocity towards the end of each reaction was fitted with the MARS software package (BMG Labtech). The rate of ATP consumption was calculated using the molar extinction coefficient of NADH (6220 M^-1^ cm^-1^). Measurements were carried out in triplicates and mean values are plotted with their associated SD.

### Thermal shift analysis

To test for correct folding of the ctMAT1 variants within the ctCAK complex, thermal shift assays were performed. CtCAK complexes were analyzed at a concentration of 2.2 mg/ml in a reaction mix containing 20 mM HEPES pH 7.5, 200 mM NaCl and 1 mM TCEP supplemented with 0.1% sypro orange (Invitrogen). Unfolding was recorded as an increase in fluorescence at an excitation wavelength of 429 nM and an emission wavelength of 610 nM using a qPCR machine (Stratagene mx 3005p). Melting temperatures represent the mean values of four replicates.

### Mass spectrometry analysis

Gel bands were excised and destained with 30% acetonitrile in 0.1 M NH_4_HCO_3_ pH 8, shrunk with 100% acetonitrile, and dried in a vacuum concentrator (Concentrator 5301, Eppendorf, Germany). Digests were performed with 0.1 μg protease (trypsin or elastase) per gel band overnight at 37 °C in 0.1 M NH_4_HCO_3_ pH 8. After removal of the supernatant, peptides were extracted from the gel slices with 5% formic acid, and extracted peptides were pooled with the supernatant.

NanoLC-MS/MS analyses were performed on an Orbitrap Fusion (Thermo Scientific) equipped with a PicoView Ion Source (New Objective) and coupled to an EASY-nLC 1000 (Thermo Scientific). Peptides were loaded on capillary columns (PicoFrit, 30 cm x 150 μm ID, New Objective) self-packed with ReproSil-Pur 120 C18-AQ, 1.9 μm (Dr. Maisch) and separated with a 30-min or 60-min linear gradient, respectively, from 3% to 40% acetonitrile and 0.1% formic acid and a flow rate of 500 nl/min.

Both MS and MS/MS scans were acquired in the Orbitrap analyzer with a resolution of 60,000 for MS scans and 15,000 for MS/MS scans. A mixed ETD/HCD method was used. HCD fragmentation was applied with 35% normalized collision energy. For ETD calibrated chargedependent ETD parameter were applied. A Top Speed data-dependent MS/MS method with a fixed cycle time of 3 s was used. Dynamic exclusion was applied with a repeat count of 1 and an exclusion duration of 10 s; singly charged precursors were excluded from selection. Minimum signal threshold for precursor selection was set to 50,000. Predictive AGC was used with AGC a target value of 2e5 for MS scans and 5e4 for MS/MS scans. EASY-IC was used for internal calibration.

The database search was performed against a custom database (about 500 proteins) containing the protein sequences of interest with the PEAKS X software (BSI, Bioinformatics Solution Inc., Canada) with the following parameters: peptide mass tolerance: 10 ppm, MS/MS mass tolerance: 0.015 Da, enzyme: “none”; variable modifications: Acetylation (Protein N-term), Oxidation (M), Carbamidomethylation (C), Pyro-glu from Q, Phosphorylation (STY). Results were filtered to 0.5% or 1% PSM-FDR, respectively, by target-decoy approach. PEAKS search results were exported for Skyline. Skyline 20.1 (skyline.maccosslab.org) was applied to determine AUC values from extracted ion chromatograms of the peptides and phosphopeptides of interest.

## Acknowledgments

We would like to thank the staff from the beamlines ID29 at the European Synchrotron Radiation Facility and MX-14 at BESSY II at the Helmholtz institute in Berlin for excellent support. We also thank Florian Sauer for providing plasmids used in this study.

## Data availability

The coordinates and structure factors for the apo or ATPγS bound ctCAK complexes have been deposited in the Protein Data Bank (PDB) under the accession code XXXX and XXXX, respectively.

