## supplement for "Structural basis for CDK7 activation by MAT1 and Cyclin H"

**Supplementary information**

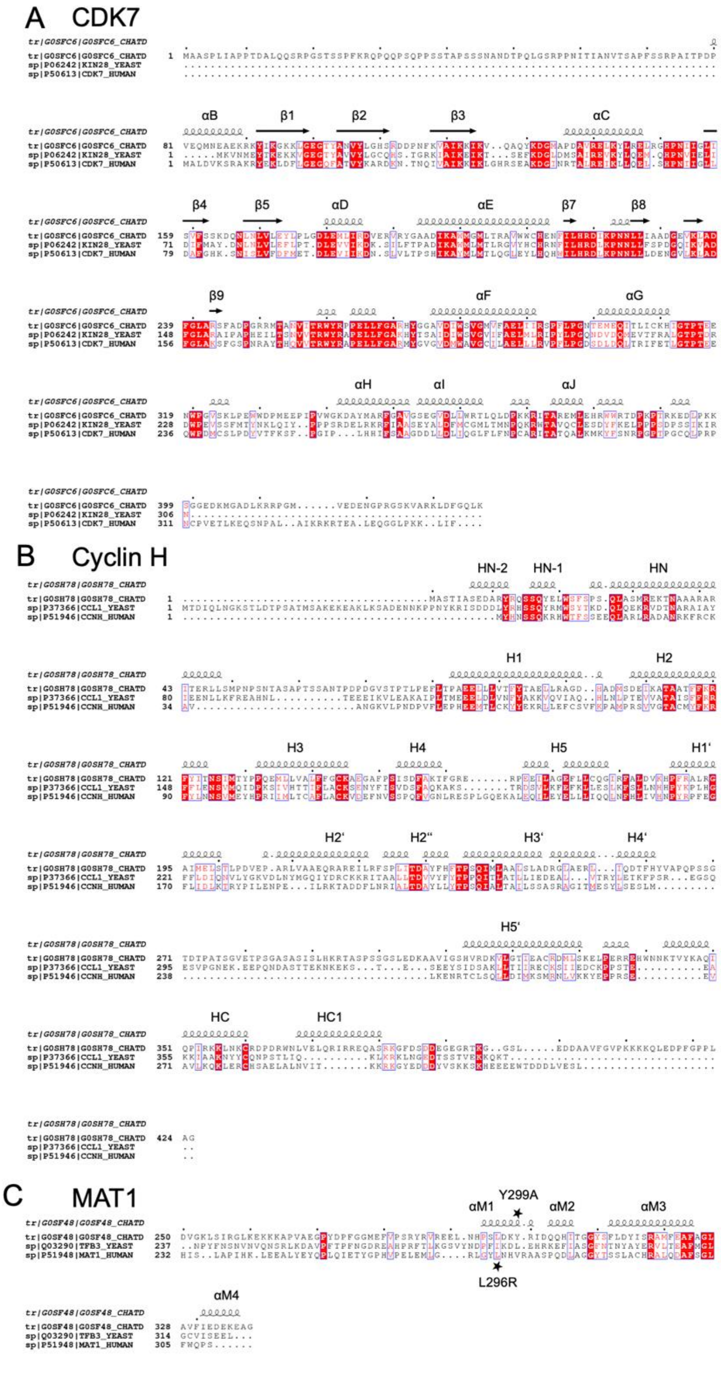


**Supplementary Figure 1:** Sequence conservation of ctCAK. **(A)** Sequence alignments of ctCDK7, Kin28 (CDK7) from *S. cerevisiae*, and human CDK7. **(B)** Sequence alignments of ctCyclin H, CCL1 (Cyclin H) from *S. cerevisiae*, and human Cyclin H. **(C)** Sequence alignments of ctMAT1short with the respective sections from, TFB3 (MAT1) from *S. cerevisiae*, and human MAT1. Secondary structure elements from ctCAKshort are depicted and labeled above the respective alignment. The alignment was prepared using the ESPRIPT server.

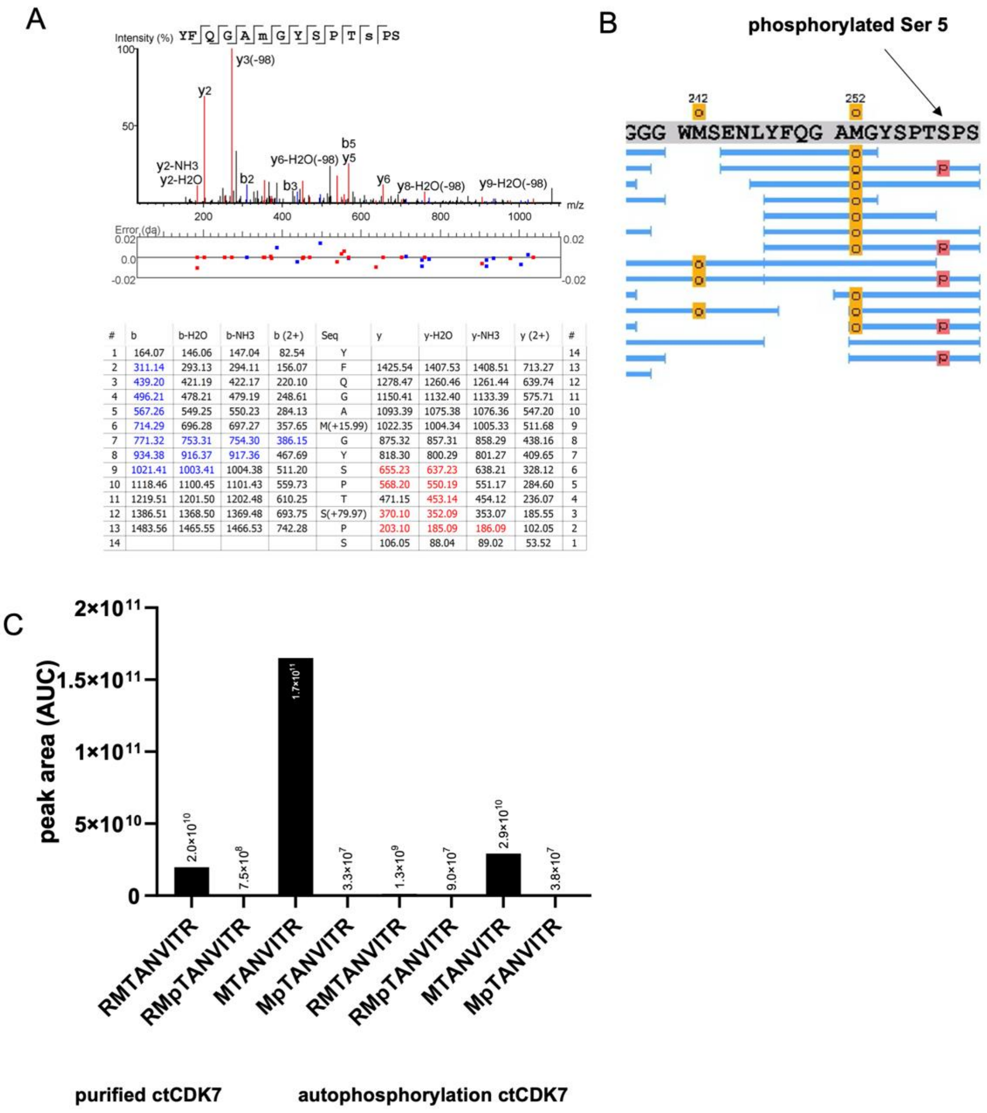


| # | b | b-H2O | b-NH3 | b (2+) | Seq | y | y-H2O | y-NH3 | y (2+) | # |
|---|---|---|---|---|---|---|---|---|---|---|
| 1 | 164.07 | 146.06 | 147.04 | 82.54 | Y | | | | | 14 |
| 2 | 311.14 | 293.13 | 294.11 | 156.07 | F | 1425.54 | 1407.53 | 1408.51 | 713.27 | 13 |
| 3 | 439.20 | 421.19 | 422.17 | 220.10 | Q | 1278.47 | 1260.46 | 1261.44 | 639.74 | 12 |
| 4 | 496.21 | 478.21 | 479.19 | 248.61 | G | 1150.41 | 1132.40 | 1133.39 | 575.71 | 11 |
| 5 | 567.26 | 549.25 | 550.23 | 284.13 | A | 1093.39 | 1075.38 | 1076.36 | 547.20 | 10 |
| 6 | 714.29 | 696.28 | 697.27 | 357.65 | M(+15.99) | 1022.35 | 1004.34 | 1005.33 | 511.68 | 9 |
| 7 | 771.32 | 753.31 | 754.30 | 386.15 | G | 875.32 | 857.31 | 858.29 | 438.16 | 8 |
| 8 | 934.38 | 916.37 | 917.36 | 467.69 | Y | 818.30 | 800.29 | 801.27 | 409.65 | 7 |
| 9 | 1021.41 | 1003.41 | 1004.38 | 511.20 | S | 655.23 | 637.23 | 638.21 | 328.12 | 6 |
| 10 | 1118.46 | 1100.45 | 1101.43 | 559.73 | P | 568.20 | 550.19 | 551.17 | 284.60 | 5 |
| 11 | 1219.51 | 1201.50 | 1202.48 | 610.25 | T | 471.15 | 453.14 | 454.12 | 236.07 | 4 |
| 12 | 1386.51 | 1368.50 | 1369.48 | 693.75 | S(+79.97) | 370.10 | 352.09 | 353.07 | 185.55 | 3 |
| 13 | 1483.56 | 1465.55 | 1466.53 | 742.28 | P | 203.10 | 185.09 | 186.09 | 102.05 | 2 |
| 14 | | | | | S | 106.05 | 88.04 | 89.02 | 53.52 | 1 |

**Supplementary Figure 2: Mass spectrometry Analysis of the phosphorylated GST-CTD peptide and T-loop phosphorylation at T253 in ctCDK7 (A)** HCD fragment ion spectrum of the precursor m/z 794.8045 (2+) corresponding to the phosphorylated substrate peptide with the sequence YFQGA(oxM)GYSPT(pS)PS generated by digest with elastase. **(B)** Closeup view of the GST-CTD sequence which was analyzed after phosphorylation by ctCAK. The position of S5 is indicated with an arrow. **(C)** Analysis of ctCDK7 directly after purification from insect cells and after addition of ATP as indicated in the Figure. The peak area (AUC) from extracted ion chromatograms (XICs) of the phosphorylated and non-phosphorylated variants of the tryptic peptides RMTANVITR and MTANVITR is shown. The Met-oxidized variant of the peptides were vastly predominant in all cases and were exclusively used for quantification.

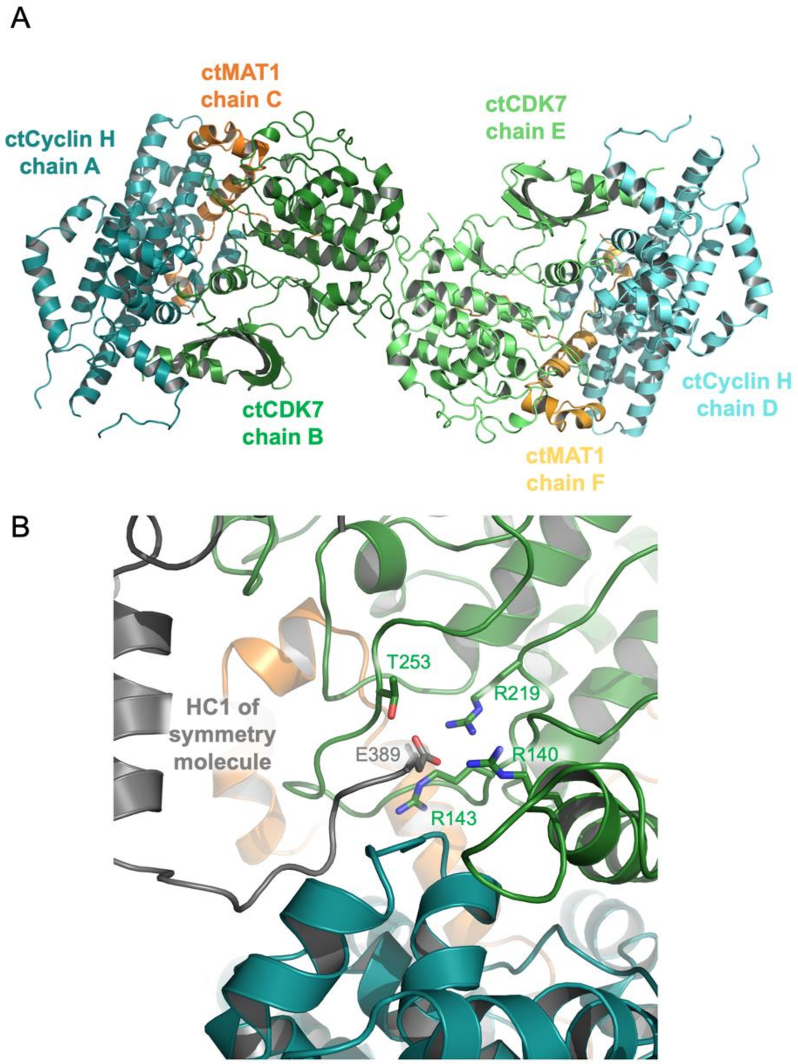


**Supplementary Figure 3:** The asymmetric unit of the ctCAK crystals and interactions of a symmetry related ctCyclin H molecule with the active site of ctCDK7 **(A)** Two ctCAK complexes are located in the asymmetric unit of the ctCAK crystals. The two heterotrimers are shown in cartoon representation with ctCyclinH in dark and light teal, ctCDK7 in dark and light green, and ctMAT1 in dark and light orange. **(B)** Closeup view of the active site of ctCDK7 with the colors as in (A). A loop from the symmetry related ctCyclin H molecule approaching the active site is shown in grey.

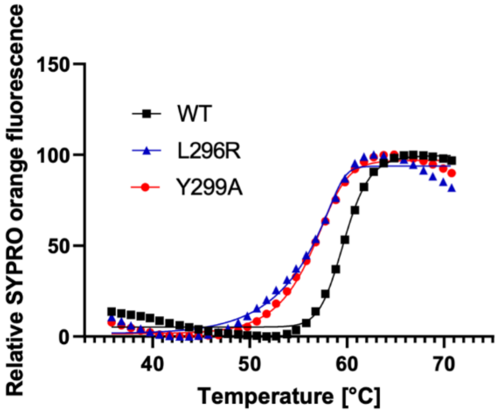


| CAK variants | mean $T_M$ [°C] |
|---|---|
| WT | 60 |
| L296R | 56 |
| Y299A | 56 |

**Supplementary Figure 4:** Thermal unfolding curves of ctCAKshort and ctCAKshort MAT1 variants and their respective melting temperatures.
